# Inhibition of AMPA receptors bound to transmembrane AMPA receptor regulatory protein γ-8 (TARP γ-8) blunts the positive reinforcing properties of alcohol and sucrose in a brain region-dependent manner

**DOI:** 10.1101/2022.12.14.520457

**Authors:** Jessica L Hoffman, Sara P Faccidomo, Seth M. Taylor, Kristina G. DeMiceli, Ashley M. May, Evan N. Smith, Ciarra M. Whindleton, Clyde W Hodge

**Author notes:** Correspondence: Clyde Hodge, Ph.D., Bowles Center for Alcohol Studies, Thurston-Bowles Building; CB#7I78, University of North Carolina at Chapel Hill, Chapel Hill, North Carolina 27599, Voice.

## Abstract

**Rationale:** The development and progression of alcohol use disorder (AUD) is widely viewed as maladaptive neuroplasticity. The transmembrane alpha-amino-3-hydroxy-5-methyl-4-isoxazole propionic acid receptor (AMPAR) regulatory protein γ8 (TARP γ-8) is a molecular mechanism of neuroplasticity that has not been evaluated in AUD or other addictions.

**Objective:** To address this gap in knowledge, we evaluated the mechanistic role of TARP γ-8 bound AMPAR activity in the basolateral amygdala (BLA) and ventral CA3 hippocampus (vHPC) in the positive reinforcing effects of alcohol, which drive repetitive alcohol use throughout the course of AUD, in C57BL/6J mice. These brain regions were selected because they exhibit high levels of TARP γ-8 expression and send glutamate projections to the nucleus accumbens (NAc), which is a key nucleus in the brain reward pathway.

**Methods and Results:** Site-specific pharmacological inhibition of AMPARs bound to TARP γ-8 in the BLA via bilateral infusion of the selective negative modulator JNJ-55511118 significantly decreased operant alcohol self-administration with no effect on sucrose self-administration in behavior-matched controls. Temporal analysis showed that reduction of alcohol-reinforced responding occurred >25 min after the onset of responding, consistent with a blunting of the positive reinforcing effects of alcohol in the absence of nonspecific behavioral effects. In contrast, inhibition of TARP γ-8 bound AMPARs in the vHPC selectively decreased sucrose self-administration with no effect on alcohol.

**Conclusions:** This study reveals a novel brain region-specific role of TARP γ-8 bound AMPARs as a molecular mechanism of the positive reinforcing effects of alcohol and non-drug rewards.

## Introduction

Alcohol use disorder (AUD) is a multiphasic neuropsychiatric condition that impacts the health and well-being of over 14-million adults each year in the United States (SAMHSA). During the binge-intoxication stage of AUD, the positive reinforcing properties of the drug promote a pattern of chronic repetitive use, followed by escalated intake over time and development of physical dependence (Koob and Volkow 2010; Wise and Koob 2013). For these reasons, research that identifies neural mechanisms that regulate the positive reinforcing properties of alcohol is crucial to understanding the etiology and progression of AUD.

The behavioral process of positive reinforcement increases the rate of adaptive behavior; however, substances of abuse can usurp neural mechanisms of reward and reinforcement and promote drug-seeking behavior. This form of maladaptive plasticity is mediated, in part, through glutamate α-amino-3-hydroxy-5-methyl-4-isoxazolepropionic acid receptor (AMPAR) activity in reward-related brain regions, including the amygdala (Kalivas 2009; Kauer and Malenka 2007; Koob 2003; 2009; Koob and Volkow 2010; Loweth et al. 2014; Marty and Spigelman 2012; McCool 2011; Roberts et al. 1996; Weiss and Koob 2001; Winder et al. 2002). Our prior research shows that CaMKII-dependent AMPAR signaling in the amygdala is required for the positive reinforcing effects of alcohol in both rats and mice (Cannady et al. 2016; Salling et al. 2016). We have also shown that AMPAR activation drives escalated alcohol self-administration and cue-induced reinstatement (Cannady et al. 2013), the latter of which is associated with increased CaMKII activity in the BLA (Salling et al. 2017). These findings indicate that CaMKII-AMPAR signaling is a neural target of alcohol that is required for the positive reinforcing effects of the drug. However, despite these and numerous other advances in understanding AMPAR regulation of alcohol drinking (reviewed by (Hopf and A. 2018)), the molecular mechanism(s) by which AMPAR activity regulates the reinforcing effects of alcohol remain to be fully elucidated.

Toward that goal, emerging evidence indicates that transmembrane AMPAR regulatory protein gamma-8 (TARP γ-8) is critical for glutamate-mediated plasticity. TARP γ-8 is a member of the calcium channel gamma subunit (*Cacng1-8;* γ1-8) family, which are the first discovered auxiliary subunits of the AMPAR or any other ligand-gated ion channel (Bissen et al. 2019). The broad family of TARPs can differentially control AMPAR pharmacology and physiology depending on cytological and neuroanatomical expression patterns (Kato et al. 2016; Kato et al. 2010). TARP γ-8 has a unique and selective anatomical expression which is highly restricted to forebrain regions such as the prefrontal cortex (PFC), hippocampus (HPC), and BLA (Fukaya et al. 2005; Maher et al. 2016). Each of these brain regions is sensitive to alcohol exposure and influences AUD-related behaviors and physiology (Hopf and A. 2018; Talani et al. 2014; White and Swartzwelder 2004). At the cellular level, TARP γ-8 is enriched in the postsynaptic density (PSD) and plays a vital role in the surface expression, trafficking, and activity of AMPARs in these brain regions (Bissen et al. 2019; Jackson and Nicoll 2011) by binding the C-terminus of GluA1 and anchoring it in the PSD (Patriarchi et al. 2018). In the forebrain, a majority of TARP γ-8 interactions with AMPARs occur at GluA1/2 heteromers (Gill et al. 2011; Herguedas et al. 2022; Schwenk et al. 2014) where the TARP γ-8 transmembrane helix 4 (TM4) associates with GluA1 and TM3 attaches between GluA1 and GluA2 to modulate both the structural and functional properties of the receptor (Herguedas et al. 2022).

This mechanistic link between TARP γ-8 and AMPAR GluA1 activity is highly relevant to understanding how alcohol may induce maladaptive plasticity during the development of AUD and other forms of addiction. GluA1-containing AMPARs are phosphorylated (e.g., activated) at serine-831 (pGluA1-S831) in the amygdala by plasticity-inducing events (Lee et al. 2013), including exposure to drugs of abuse (Conrad et al. 2008; Mameli et al. 2011; Wolf and Tseng 2012). We have shown that alcohol self-administration increases pGluA1-S831 expression and synaptic insertion of calcium-permeable GluA2-lacking AMPARs (CP-AMPARs) in the BLA, and that membrane insertion of GluA1-containing AMPARs is required for the positive reinforcing effects of alcohol (Faccidomo et al. 2021). Moreover, we recently found that systemic pharmacological inhibition of TARP γ-8 bound AMPARs significantly reduced operant alcohol self-administration by male, but not female, C57BL/6J mice (Hoffman et al. 2021), which was the first evidence that TARP γ-8 mediates any form of substance abuse. Together, these findings suggest the novel hypothesis that TARP γ-8 regulates the positive reinforcing properties of alcohol via modulation of AMPAR activity in specific reward-related brain regions.

To address this question, the present experiments were designed to determine the effects of reversible pharmacological inhibition of TARP γ-8 bound AMPARs in the BLA and vHPC on operant alcohol self-administration by male C57BL/6J mice. We focused on the BLA and vHPC as potential sites of action based on high levels of TARP γ-8 expression (Maher et al. 2017; Maher et al. 2016) and our prior work demonstrating that glutamate activity in these regions regulates alcohol-related behavioral pathologies (Cannady et al. 2016; Faccidomo et al. 2021; Faccidomo et al. 2019; Salling et al. 2016; Salling et al. 2017; Spanos et al. 2012). C57BL/6J mice were trained to self-administer alcohol in operant conditioning chambers using our well-characterized method that compares responding reinforced by sweetened alcohol to parallel behavior-matched sucrose-only controls (Faccidomo et al. 2009; Faccidomo et al. 2019; Faccidomo et al. 2016b; Faccidomo et al. 2015; Faccidomo et al. 2018; Salling et al. 2016). To assess mechanistic regulation of self-administration behavior, the selective inhibitor of TARP γ-8 bound AMPARs, JNJ-55511118 (JNJ-5), was microinjected in the BLA or vHPC prior to operant self-administration sessions. JNJ-5 is a high affinity negative modulator of GluA1 containing AMPARs bound to TARP γ-8 that disrupts the interaction between TARP γ-8 and AMPAR GluA subunits, effectively inhibiting postsynaptic AMPAR activity (Maher et al. 2016); this selective disruption results in strong dose-dependent inhibition of AMPAR-mediated transmission and anticonvulsant properties in rodent models (Maher et al. 2016). Following self-administration studies, an effective dose of JNJ-5 was also evaluated for potential nonspecific effects on locomotor and anxiety-like (thigmotaxis) behavior.

This study is the first to examine TARP γ-8 as a driving force of the reinforcing effects of any drug of abuse via activity within a brain reward pathway and provides further support for the premise that targeting this AMPAR subclass may be a viable strategy for developing medications to treat behavioral pathologies associated with AUD (Hoffman et al. 2021) or other neurological conditions (Maher et al. 2017).

## Materials and methods

### Animals

Male C57BL/6J mice (The Jackson Laboratory, Bar Harbor, ME USA) arrived in our colony room at 8-10 weeks old and habituated to the environment for 1 week. Mice were group-housed in Techniplast cages (28 x 17 x 14 cm) containing a plastic hut and nestlet for environmental enrichment. Food (Purina chow) and water were available ad libidum unless otherwise indicated. The colony and behavioral testing rooms were temperature (21±1°C) and humidity (40±2%) controlled and maintained on a 12h:12h reverse light/dark cycle (dark at 0700). All experimental procedures involving mice were approved by the Institutional Care and Use Committee at the University of North Carolina at Chapel Hill and conducted as recommended by the Guide for the Care and Use of Laboratory Animals (National Research Council (U.S.). Committee for the Update of the Guide for the Care and Use of Laboratory Animals. et al. 2011).

### Drugs

#### Reinforcing solutions

Sweetened alcohol (alcohol 9% v/v + sucrose 2% w/v) was diluted from a 95% ethanol stock (Pharmco Products Inc., Brookfield, CT, USA) to 9% v/v with distilled water and sweetened with sucrose (2% w/v). The sucrose only solution (2% w/v) was also prepared with sucrose and distilled water.

#### JNJ-55511118

5-[2-Chloro-6-(trifluoromethoxy)phenyl]-1,3-dihydro-2H-benzimidazol-2-one (JNJ-5; Tocris, Minneapolis, MN) is a high affinity, negative modulator of TARP γ-8 bound AMPA receptors(Maher et al. 2016). JNJ-5 (0.0, 0.3, 1.0, and 2.0ug/ul) was suspended in aCSF and bilaterally microinfused into target brain regions.

#### Apparatus

Operant chambers (Med-Associates, St. Albans, VT) were housed in sound-attenuating cubicles to reduce extraneous noise. Chambers were computer-interfaced for automated control of inputs (e.g., recording of mouse behavior) and outputs (e.g., delivery of solutions) using commercially available software (MED-PC for Windows v5.0). Each chamber was equipped with two ultra-sensitive retractable response levers, positioned on opposite walls below a cue light. Levers were programmed to be either “active” (reinforced) or “inactive” and responses were recorded for both. A syringe pump was connected to a drinking trough positioned in the center of the chamber and adjacent to the active lever. Reinforced, active lever presses were accompanied by secondary cues including a cue light (800ms) and the pump sound. Responses during pump run time were recorded but produced no programmed consequences.

#### Homecage Exposure to Reinforcing Solutions

After habituation to the colony room (1 week), two drinking bottles were attached to each mouse homecage; one bottle contained sweetened alcohol (9% v/v + sucrose 2% w/v) or sucrose only (2% w/v) and the other bottle contained water. Mice were able to freely consume the reinforcing solution vs. water for two weeks prior to operant training reducing neophobic responses to the solutions during operant training (Faccidomo et al. 2015).

### Procedural sequence

The following sections describe the experimental procedures shown in Figure 1.

**Figure 1.**
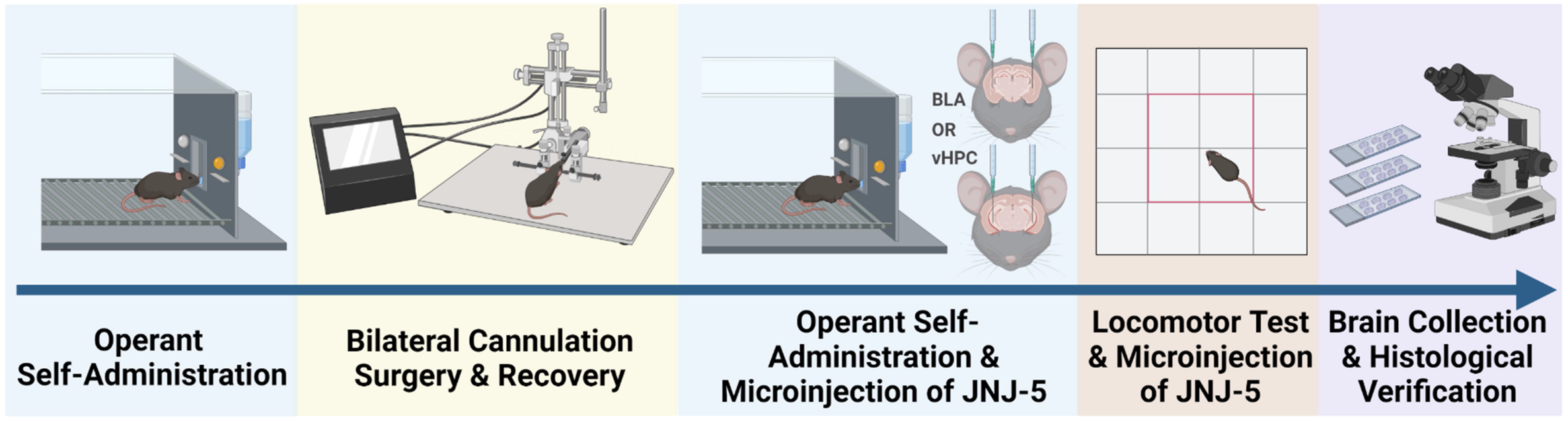
Sequence of experimental procedures. Mice were first trained to self-administer sweetened alcohol or sucrose-only in operant conditioning chambers. Training was followed by bilateral cannulation surgery and recovery and then a return to operant responding. Mice were then microinjected with JNJ-5 (0, .3, 1, 2ug/ul) into the BLA or vHPC prior to operant self-administration to determine the effects of inhibiting TARP γ-8 bound AMPAR on alcohol or sucrose self-administration. The JNJ-5 microinjections were repeated prior to an open field test to account for any non-specific locomotor or anxiety-like (thigmotaxis) effects. Finally, brains were removed for histological verification of cannula placement.

### Operant Self-Administration

#### Training and acquisition

Training and acquisition of mouse operant self-administration of sweetened alcohol (9% v/v + sucrose 2% w/v) or sucrose (2% w/v) in separate groups of mice was conducted as described previously (Faccidomo et al. 2009; Hoffman et al. 2021; Salling et al. 2008). Briefly, initial training occurred during 3-4 consecutive overnight (16hr) sessions; the number of sessions was determined by learning contingent lever pressing. Mice were water restricted 24hrs prior to the initial training session. Initial lever press responses were reinforced on a fixed ratio 1 (FR1) schedule that increased to FR2, FR3, and to a final value of FR4 over the training session(s) with each increment of 25 reinforcers earned at each FR value. Active lever presses associated with a reinforcer activated the syringe pump to deliver 0.014ml of the reinforcing solution into the drinking trough. After completing the session, an experimenter confirmed fluid intake via visual inspection of the drinking trough. All subsequent operant self-administration sessions took place during the dark phase of the light/dark cycle (between 1200-1600) and lasted for 60min and were on a FR4 schedule. These 1hr daily sessions were conducted M-F unless otherwise noted. Our previous work using this protocol has resulted in pharmacologically relevant blood alcohol content (BAC) (Faccidomo et al. 2009) levels though blood samples were not taken during this study to minimize behavioral disruption and possible stress-induced alterations of AMPARs (Bats et al. 2013; Kuniishi et al. 2020).

### Surgery

After a baseline of 40-42 operant self-administration sessions, mice underwent surgery for bilateral cannula implantation. Mice were anesthetized with vaporized isoflurane (1.5-2.5%) and placed into a stereotaxic frame (Kopf Instruments, Tujunga, CA). Hair was removed from the scalp, and the surface was scrubbed with ethanol and beta iodine to sterilize the skin prior to incision. Microclips were used to pull back skin to expose the skull to identify bregma. Using a digital arm, holes were drilled bilaterally at the following coordinates derived from Paxinos and Franklin’s Mouse Brain Atlas, (Franklin and Paxinos 2019): BLA: AP=-1.2 and ML=+/-3.3 from bregma, DV=-2.6mm from dura; vHPC: AP=-3.0 and ML=+/-3.3 from bregma, DV=-2.0mm from dura. Next, 26-gauge guide cannula (Plastics One, Roanoke, VA) were implanted positioned 2mm above either the BLA or vHPC and cemented to the skull with dental cement (Durelon, Butler Schein, Dublin, OH). A 33-gauge obturator extending .5mm beyond the cannula tip was inserted into the guide cannula after surgery and moved daily to prevent blockage and scarring. Mice were monitored and kept warm until awakening from anesthesia, returned to the colony room, and administered ibuprofen for acute pain. A bitter tasting deterrent solution was brushed onto the head-mount and obturator to discourage damage from cage-mates. After a 7-day recovery period, mice returned to operant self-administration.

### Microinjections

Site-specific microinjections were conducted as previously reported (Besheer et al. 2012; Faccidomo et al. 2016a; Hodge et al. 1996; Samson and Hodge 1993; Schroeder et al. 2003). After re-establishing baseline self-administration, mice were habituated to the microinjection procedure with sham injections, which involved insertion of the injector and running the pump for 4min (not connected to the injector). After the sham injection, mice were immediately placed into the operant chamber for the 60min session. Once habituated, separate groups of mice were administered JNJ-5 (0.0, 0.3, 1.0 and 2.0μg/.5μl/side) via bilateral microinjection in the BLA or vHPC according to a randomized Latin-square dosing design with a minimum of 3 days between injections. Mice were unrestrained during the JNJ-5 infusions; the infusions lasted for 4min at a rate of 0.125μl/min/side using a 1ul Hamilton syringe connected to a syringe pump (Harvard Apparatus, Holliston, MA). After the infusion, the injector remained in place for 1min to allow drug diffusion and minimize vertical capillary action. Mice were then immediately placed into the operant chamber for the 60min self-administration session.

### Locomotor Activity

Locomotor testing was conducted to evaluate potential nonspecific motor or anxiety-like effects of JNJ-5 infusion (Agoglia et al. 2016; Agoglia et al. 2015; Besheer et al. 2006; Faccidomo et al. 2015; Hodge et al. 2002; Kelley et al. 2003). Mice were habituated to open field chambers (Med-Associates) for 2hrs, one week prior to locomotor testing. Microinjections of JNJ-5 (0.0, 0.3, 1.0 and 2.0 μg/side; counterbalanced design) into the BLA or vHPC were conducted immediately prior to a 60min locomotor activity assessment. Distance traveled (cm) was computer-recorded every 100ms in the open field chamber via two sets (x and y axes) of 16 pulse-modulated infrared photobeams. Thigmotaxis (distance in periphery vs center) was derived as a measure of anxiety-like behavior. Locomotor activity assessments occurred at least 4 days apart.

### Histological Verification

After completion of all microinjections (prior to operant self-administration and locomotor activity), mice were deeply anesthetized with sodium pentobarbital and intracardially perfused with 0.9% phosphate buffered saline (PBS) and 4% paraformaldehyde (PFA). Brains were extracted, coronally sliced (50um) on a vibratome (Lecia VT100S, Leica Biosystems, Wetzlar, Germany), and stained with cresyl violet for histological verification. Data were used only from mice verified to have received bilateral infusion in the BLA or vHPC.

### Statistical Analysis

All analyses were conducted with Prism 9 (Graphpad). **Repeated-measures (RM) ANOVA or mixed-effects model (REML) analyses were** used to compare groups with corrected multiple comparisons (Holm-Šídák) **when appropriate**.

## Results

### EtOH reinforcement is regulated by TARP γ-8 bound AMPARs in the BLA

To determine if the activity of TARP γ-8 bound AMPARs in the BLA is required for EtOH reinforcement, the selective inhibitor JNJ-5 (0–2.0 μg/.5μl/side) was microinjected in the BLA prior to operant self-administration sessions in C57BL/6J mice (Fig 2A). RM-ANOVA showed that intra-BLA infusion of JNJ-5 selectively decreased total EtOH-reinforced responding, *F*(3,21)=5.481, *p*=0.01. Follow up analysis with Holm-Šídák’s multiple comparison tests show that all doses of JNJ-5 decreased total EtOH-reinforced responses (Fig 2B). Reduced total EtOH-reinforced responses was associated with a significant decrease in EtOH intake (g/kg), *F*(3,21)=3.564, *p*=0.03, which was a function of decreases following JNJ-5 (0.3 and 2.0 μg/.5μl/side) doses (Fig 2B). RM-ANOVA showed no differences in inactive lever responding, *F*(3,21)=0.111, *p*=0.95, or total headpokes/reinforcer, *F*(3,21)=2.519, *p*=0.086 for mice consuming EtOH (Fig 2B), suggesting that the reduction in EtOH-reinforced responding was not associated with nonspecific motor or consummatory effects, respectively.

**Figure 2.**
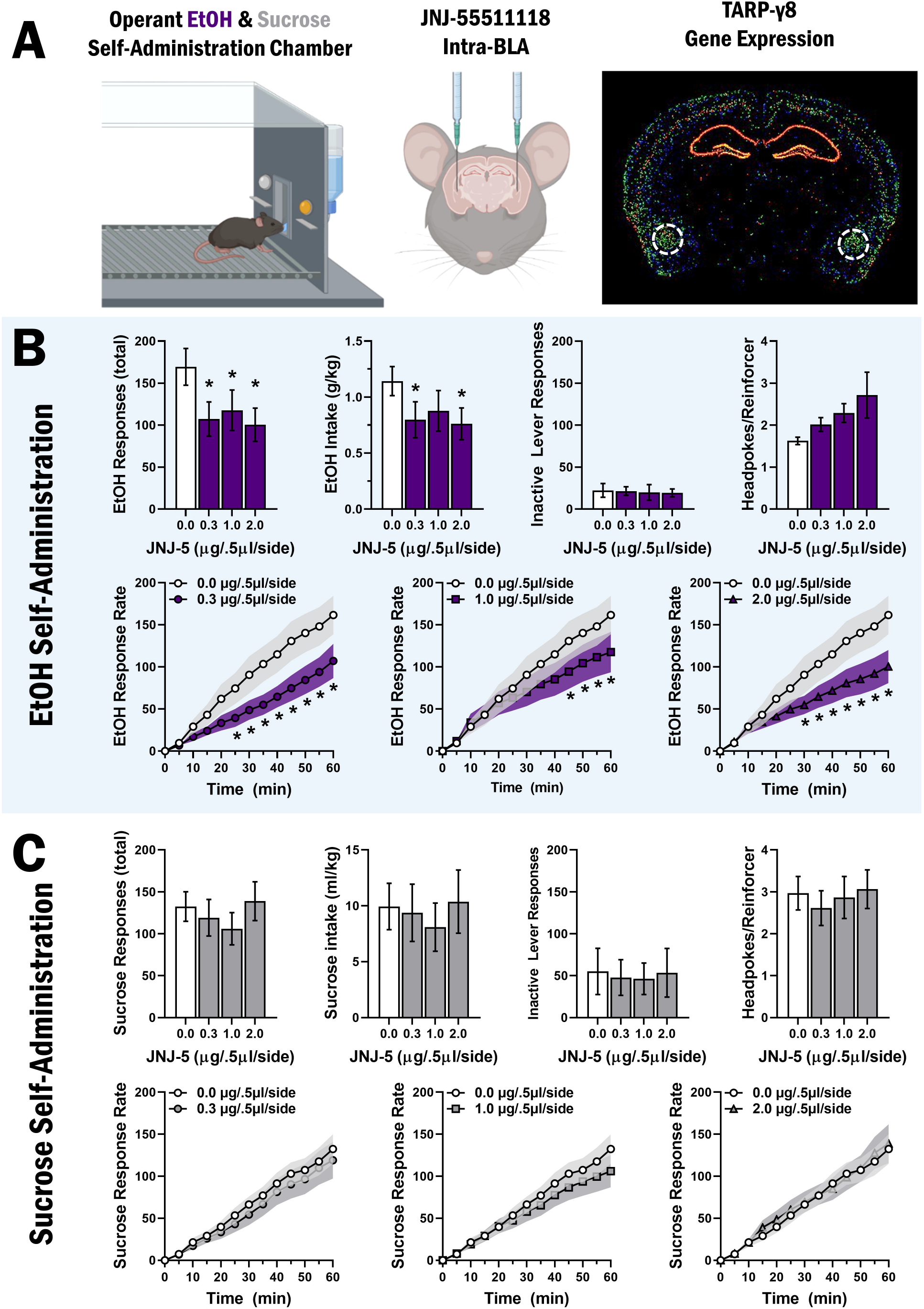
Inhibition of TARP v-8 bound AMPARs in the BLA selectively decreases EtOH reinforced responding. (**A**) Experimental schematic for intra-BLA microinjections of JNJ-55511118 prior to operant selfadministration of alcohol (n=8) and sucrose-only (n=9), and BLA TARP γ-8 gene expression denoted by dashed white lines, Allen Brain Atlas. (**B-C**) Bar graphs (top rows) show MEAN±SEM parameters of operant ethanol (**B**) and sucrose (**C**) self-administration plotted as a function of dosage of JNJ-5 infused in the BLA. Line graphs (bottom rows) show MEAN±SEM rate of ethanol (**B**) and sucrose (**C**) reinforced responding (cumulative responses / 5-min interval) as compared to vehicle control for each dose of JNJ-5 tested. *denotes statistically significant difference as compared to vehicle, p<.05.

To directly evaluate regulation of EtOH reinforcer function, we also examined the impact of intra-BLA infusion of JNJ-5 on EtOH-reinforced response rate. A mixed-effects model analysis of EtOH-reinforced response rate revealed a Time x Dose interaction, *F*(33,231)=5.156, *p*<0.0001, and main effects of Time, *F*(11,77)=30.68, *p*<.0001, and Dose, *F*(3,21)=3.722, p=0.03. Post hoc Holm-Šídák’s multiple comparisons reveal statistically significant differences between JNJ-5 0.0 μg/.5μl/side and .3 μg/.5μl/side doses from 25-60min, additional differences between 0.0 μg/.5μl/side and 2. μg/.5μl/side from 30-60min, and additional differences between 0. μg/.5μl/side and 1.0 μg/.5μl/side from 45-60min (**Fig 2B, bottom row**).

### TARP γ-8 bound AMPAR inhibition in the BLA has no effect on sucrose reinforcement

To assess EtOH-reinforcer specificity, we inhibited TARP γ-8 bound AMPARs in the BLA in behavior-matched controls trained to self-administer sucrose-only (2% w/v). Intra-BLA infusion of JNJ-5 (0-2 μg/.5μl/side) has no effect on sucrose reinforced responses *F*(3,23)=1.143, *p*=0.35 or on sucrose intake (ml/kg), *F*(3,23)=0.599, *p*=0.62 (**Fig 2C**). Mixed-effects model analysis shows no differences in inactive lever responding, *F(*3,23)=0.364*, p=0.78*, or headpokes/reinforcer, *F*(3,23)=1.915, *p*=0.15 (**Fig 2C**) indicating that JNJ-5 administered in the BLA had no effect on non-drug reinforcement, motor performance, or consummatory behavior. Mixed-effects model analysis of sucrose-reinforced response shows a main effect of time, *F*(11,88)=40.88, *p*<.0001 (**Fig 2C, bottom row**), indicating no effect of JNJ-5 in the dose range tested.

### TARP γ-8 bound AMPAR inhibition in vHPC has no effect on EtOH reinforcement

To evaluate brain regional specificity of TARP γ-8 regulation of EtOH reinforcement, JNJ-5 was microinjected in vHPC prior to operant self-administration sessions in C57BL/6J mice (**Fig 3A**). In contrast to the BLA, mixed-effects model analysis shows that Intra-vHPC JNJ-5 infusion has no effect on total EtOH-reinforced responding, *F*(3,20)=0.1614, *p*=0.92(**Fig 3B**). Similarly, RM-ANOVA showed no effect on EtOH intake (g/kg), *F*(3,20)=0.055, *p*=0.982 (**Fig 3B**). Mixed-effects model shows no differences in inactive lever responding, *F*(3,20)=0.3407, *p*=0.80, or headpokes/reinforcer, *F*(3,20)=1.028, *p*=0.40, for mice consuming EtOH (**Fig 3B**). An additional mixed-effects model analysis of the rate of EtOH-reinforced response rate shows a main effect of Time, *F*(11,99)=55.70, *p*<.0001(**Fig 3B**), indicating no effect of JNJ-5.

**Figure 3.**
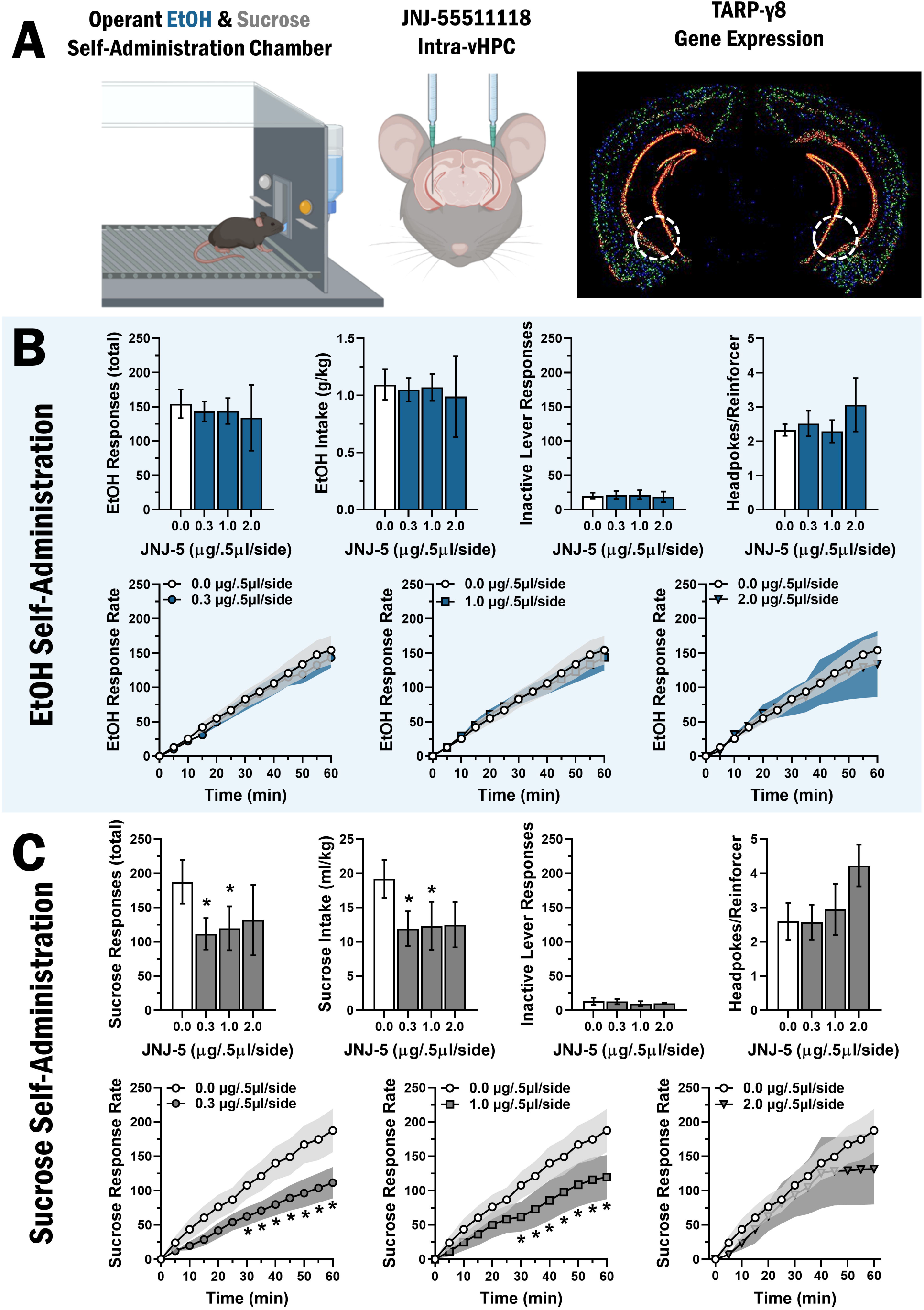
TARP v-8 bound AMPAR inhibition in vHPC selectively decreases sucrose responding. (**A**) Experimental schematic for intra-vHPC microinjections of JNJ-55511118 prior to operant self-administration of alcohol (n=8) and sucrose-only (n=9), and vHPC TARP γ-8 gene expression denoted by dashed white lines, Allen Brain Atlas. (**B-C**) Bar graphs (top rows) show MEAN±SEM parameters of operant ethanol (**B**) and sucrose (**C**) self-administration plotted as a function of dosage of JNJ-5 infused in the vHPC. Line graphs (bottom rows) show MEAN±SEM rate of ethanol (**B**) and sucrose (**C**) reinforced responding (cumulative responses / 5-min interval) as compared to vehicle control for each dose of JNJ-5 tested. *denotes statistically significant difference as compared to vehicle, p<.05.

### TARP γ-8 bound AMPAR inhibition in vHPC decreases sucrose-reinforced responding

We also sought to determine potential regulation of sucrose reinforcement by TARP γ-8 bound AMPARs in the vHPC. Results showed that intra-vHPC infusion of JNJ-5 (0-2 μg/.5μl/side) decreased sucrose-reinforced responding, *F*(3,14)=4.288, *p*=0.02 and intake (ml/kg), *F*(3,14)=5.856, *p*=0.01 (**Fig 3C**). Mixed-effects model analysis shows no differences in inactive lever responding, *F*(3,14)=0.1173, *p*=0.95, and no difference in headpokes/reinforcer, *F*(3,14)=0.5789, *p*=0.64 (**Fig 3C**).

Evaluation of sucrose-reinforced response rate by mixed-model analysis shows a Time x Dose interaction *F*(33,147)=2.413, *p*<0.0001, and main effects of Time, *F*(11,77)=34.84, *p*<.0001, and Dose, *F*(3,21)=5.407, *p*<0.01. Post hoc Holm-Šídák’s multiple comparisons revealed statistically significant differences between JNJ-5 0.0μg/.5μl/side and 0.3 μg/.5μl/side doses from 25-60min, additional differences between 0.0μg/.5μl/side and 2.0 μg/.5μl/side from 30-60min, and additional differences between 0.0 μg/.5μl/side and 1.0 μg/.5μl/side from 45-60min (**Fig 3C, bottom row**).

### Site-Specific TARP γ-8 bound AMPAR inhibition does not alter locomotor activity or thigmotaxis

To determine if decreases in operant self-administration induced by site-specific inhibition of TARP γ-8 bound AMPAR activity are due to changes in motor function, additional microinjections of JNJ-5 into the BLA (**Fig 4A-C**) and vHPC (**Fig 4D-F**) were performed prior to tests of open-field locomotor activity. RM-ANOVA shows no difference in distance traveled after BLA microinjection of JNJ-5 in mice exposed to EtOH, *F*(3,24)=2.728, *p*=0.07, or sucrose, *F*(3,27)=0.04578, *p*=0.99 (**Fig 4B**). Similarly, mixed-effects model analysis of % distance travelled in center of an open field following BLA JNJ-5 microinjection showed no differences in mice exposed to EtOH, *F*(3,31)=0.1254, *p*=0.94, or sucrose, *F*(3,35)=0.1348, *p*=0.94 (**Fig 4C**). Likewise, Mixed-effects model analysis of vHPC microinjection of JNJ-5 showed no differences in distance traveled in mice exposed to EtOH, *F*(3,13)=2.951, *p*=0.07, or sucrose, *F*(3,13)=0.9423, *p*=0.45 (**Fig 4E**). Finally, mixed-effects model analysis also showed no effect of vHPC microinjection of JNJ-5 in % distance traveled in the center of an open field for mice exposed to EtOH, *F*(3,13)=0.2354, *p*=0.87, or sucrose, *F*(3,13)=1.402, *p*=0.30 (**Fig 4F**).

**Figure 4.**
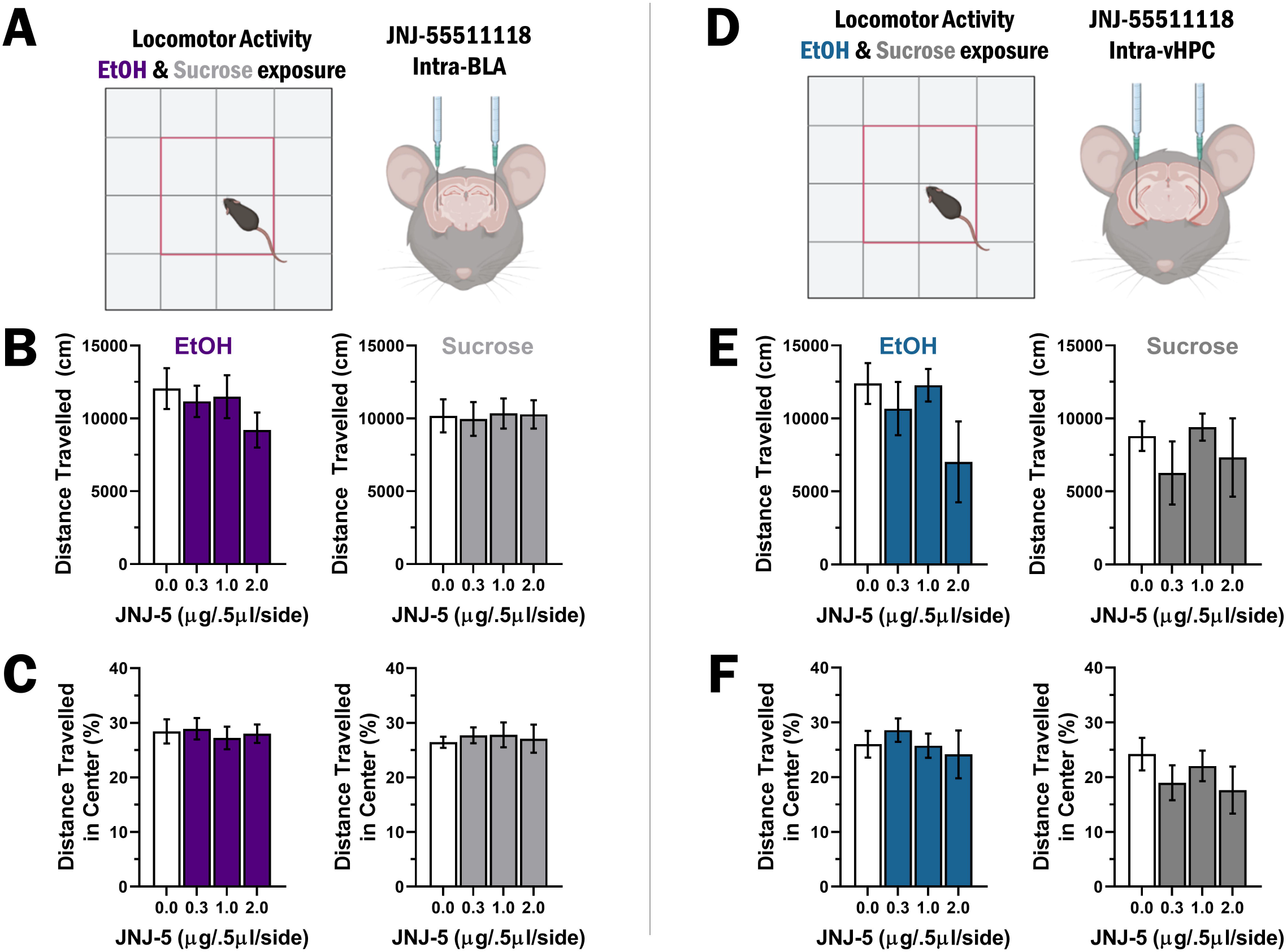
Site-Specific inhibition of TARP v-8 bound AMPAR has no effect on locomotor activity or anxiety-like behavior. (**A**) Experimental schematic showing open-field with center zone and intra-BLA microinjections prior to open field test. (**B**) Motor activity shown as mean ± SEM distance traveled (cm) during each 60min session for EtOH (left, blue bars, n=9) and sucrose (right, gray bars, n=10) exposed mice plotted as a function of JNJ-5 dosage in the BLA. (**C**) Thigmotaxis shown as mean ± SEM distance traveled (cm) in the center of the open field for EtOH (left, blue bars) and sucrose (right, gray bars) exposed mice plotted as a function of JNJ-5 dosage in the BLA. (**D**) Experimental schematic of the apparatus and intra-vHPC microinjections prior to open-field test. (**E**) Motor activity plotted as Mean±SEM distance traveled (cm) during 60min sessions for EtOH (left, blue bars, n=9) and sucrose (right, gray bars, n=9) exposed mice plotted as a function of JNJ-5 dosage in the VHPC. (**F**) Thigmotaxis shown as Mean±SEM distance traveled (cm) in the center of the open field for EtOH (left, blue bars) and sucrose (right, gray bars) exposed mice plotted as a function of JNJ-5 dosage in the vHPC.

## Discussion

Our prior research shows that inhibition of TARP γ-8 bound AMPARs via systemic administration of the selective inhibitor JNJ-5 significantly decreases operant alcohol self-administration in C57BL/6J mice (Hoffman et al. 2021). The present study extends this finding and identifies TARP γ-8 in the BLA as a novel site-specific molecular mechanism of the positive reinforcing effects of alcohol. Here we show that microinjection of JNJ-5 (0 – 2μg/.5μl/side), a high affinity negative modulator of TARP γ-8 bound AMPARs, in the BLA significantly reduced the total amount and rate of operant alcohol (sweetened) self-administration by C57BL/6J mice. By contrast, JNJ-5 infusion in the BLA had no effect on sucrose-only self-administration in parallel behavior-matched controls, which suggests alcohol reinforcer-specificity. Critical control measures within the operant procedure and separate tests found no disruption of motor, consummatory, or anxiety-like behavior. This selective reduction in alcohol-reinforced response rate indicates that TARP γ-8 modulated AMPAR activity in the BLA is required for the full expression of the positive reinforcing properties of alcohol.

The BLA plays a prominent role in reinforcement processes through glutamatergic projections to the NAc (Wright et al. 1996), which are both necessary and sufficient to promote reward-seeking behavior (Stuber et al. 2011). We have shown previously that operant alcohol self-administration increases postsynaptic insertion of GluA1-containing AMPARs in BLA neurons that project to the NAc, and that this membrane trafficking of GluA1 in the BLA is required for the reinforcing effects of alcohol (Faccidomo et al. 2021). TARP γ-8 modulates postsynaptic glutamate signaling and behavioral plasticity by binding the AMPAR GluA1 subunit C-terminus and anchoring it in the PSD (Park et al. 2016; Patriarchi et al. 2018). Thus, TARP γ-8 may play a role in alcohol-induced synaptic insertion of GluA1-containing AMPARs in BLA projection neurons and in alcohol reinforcement processes via modulation of excitatory projections from the BLA to reward-related neural structures including the NAc.

Although the present results show a significant role for TARP γ-8 in the BLA, they also suggest that TARP γ-8 modulation of AMPAR activity in the vHPC does not regulate the reinforcing properties of alcohol. Site-specific infusion of JNJ-5 in the vHPC had no effect on any measure of operant alcohol self-administration, or measures of motor activity and anxiety-like behavior. This finding was surprising as TARP γ-8 is highly abundant in the projection regions of the hippocampus (Maher et al. 2016) where CaMKII and TARP γ-8 have been shown to regulate synaptic and behavioral plasticity (Park et al. 2016) and alcohol enhances glutamatergic output from the vHPC to NAc neurons expressing dopamine D1 receptors (Kircher et al. 2019), which are known to regulate alcohol self-administration (Hodge et al. 1997). Moreover, the hippocampus and NAc interactively regulate the glutamatergic component of alcohol’s discriminative stimulus properties (Hodge and Cox 1998), which are fundamental to reinforcement processes (Stolerman 1992). Thus, these results are consistent with the conclusion that there is brain region-specific regulation of alcohol reinforcement by AMPARs associated with TARP γ-8 in the BLA. However, since TARP γ-8 and AMPARs are expressed throughout the hippocampus and various cortical areas, further research is warranted to assess this hypothesis.

A key finding from this study is that the positive reinforcing properties of alcohol and those of non-drug rewards, such as sucrose, may be regulated by TARP γ-8 bound AMPARs in a brain region-dependent manner. We found that JNJ-5 infusion in the vHPC inhibited sucrose-reinforced responses with no effect on alcohol, whereas an opposite alcohol-specific effect was observed in the BLA. Although the mechanism for this double dissociation is unclear, one plausible hypothesis is that alcohol exposure may differentially alter TARP γ-8 or AMPAR expression in a manner that alters response to JNJ-5. As noted above, our prior work shows that alcohol self-administration upregulates both CP-AMPAR synaptic expression and GluA1 activity (GluA1-S831 phosphorylation) in the BLA to a greater extent than sucrose and, in turn, GluA1 membrane insertion is required for the positive reinforcing effects of alcohol (Faccidomo et al. 2021). Thus, if alcohol selectively upregulates TARP γ-8 expression or activity in the BLA, this may increase the population of TARP γ-8 bound AMPARs in the BLA and promote behavioral response to JNJ-5 via enhanced output to downstream projection regions. By contrast, chronic alcohol exposure reduces neuronal excitability (Bach et al. 2021) and GluA1 expression (Yao et al. 2021) in the vHPC, suggesting that a potential downregulation of TARP γ-8 by alcohol might decrease the population of TARP γ-8 bound AMPARs and blunt response to JNJ-5. Interestingly, our observation that inhibition of TARP γ-8 bound AMPARs in the vHPC decreases the reinforcing effects of sucrose is consistent with evidence showing that glutamate neurotransmission in the vHPC regulates feeding (Kanoski and Grill 2017) and, specifically, that sucrose intake increases total AMPAR GluA1 subunit expression in the vHPC (Ross et al. 2019). To address these questions, future studies can evaluate the impact of alcohol and/or sucrose on brain regional TARP γ-8 gene and protein expression and assess a broader dose range of JNJ-5 in each brain region to account for potential differential sensitivity.

It is widely recognized that AMPAR antagonists produce significant side-effects including locomotor and cognitive deficits. However, site-specific inhibition of TARP γ-8 bound AMPARs was without effect on motor activity or thigmotaxis (anxiety-like behavior). Moreover, lack of significant reductions in operant behavior during the initial onset of responding, or on reinforcer consumption (headpokes), suggests lack of memory deficits. This is consistent with our prior observation that systemic administration of JNJ-5 reduced alcohol self-administration by male, but not female, mice in the absence of motor effects (Hoffman et al. 2021). Thus, blocking only the subclass of AMPARs associated with TARP γ-8 may provide therapeutic efficacy in the absence of negative side-effects; however, further work needs to explore the possibility of sexually dimorphic sensitivity to JNJ-5 (Hoffman et al. 2021) and the potential nonspecific impact of higher doses in females.

In conclusion, discovering mechanistically driven therapeutic interventions for behavioral pathologies associated with AUD remains a challenge. Despite a plethora of preclinical and clinical evidence identifying glutamate neurotransmission as a potential target for alcohol medications (Heilig and Egli 2006; Holmes et al. 2013), selective treatment approaches remain to be developed. Toward that goal, use of AMPAR negative modulators is appealing for pharmacological treatment of AUD and other CNS disorders involving increased neuronal excitability. However, nonspecific inhibition of AMPARs produces undesirable side effects including sedation and memory disruption (Rogawski 2011). Thus, there is a need for novel AMPAR-targeted therapeutic strategies that selectively modulate disease-specific brain regions or pathways. Given the restricted anatomical expression of TARP γ-8 (e.g., hippocampus, frontal cortex, amygdala), this AMPAR auxiliary protein has been proposed as a novel target for modulating excitability in specific brain regions via systemic treatment (Gill and Bredt 2011; Maher et al. 2017). In support of TARP γ-8 as a therapeutic target for AUD, we showed previously that systemic administration of JNJ-5 selectively reduces the reinforcing effects of alcohol in mice (Hoffman et al. 2021) and, in the present study, localized this therapeutic-like effect to the BLA. This suggests that systemic treatment with JNJ-5 reduces the reinforcing effects of alcohol by inhibition of TARP γ-8 associated AMPARs selectively within the amygdala, a key component of the brain’s reward pathway (Koob 1999; 2003). It will be important to determine if TARP γ-8 regulates other behavioral pathologies including relapse-like behavior and dependence-induced escalated alcohol use, both of which characterize advanced stages of AUD and lack treatment approaches.

## Acknowledgements and Disclosures

Research reported in this publication was supported by the National Institute on Alcohol Abuse and Alcoholism of the National Institutes of Health under award numbers R01 AA028782 (CWH), P60 AA011605 (CWH), and F32 AA028993 (JLH), and by the Bowles Center for Alcohol Studies at The University of North Carolina at Chapel Hill.

JLH, SF, and CWH designed the experiments. JLH, SF, and KGD performed the surgeries. JLH, SMT, KGD, AMM, and ENS performed the experiments. JLH and CWH analyzed the data. JLH, SF, and CWH interpreted the data, wrote, and edited the manuscript.

The authors wish to thank our dedicated team of undergraduate research assistants and research technicians for their assistance. Specifically, we thank Eric Homberger, Julie Lee, and LC Wong for their help with behavioral tasks and laboratory upkeep.

The authors report no biomedical financial interests or potential conflicts of interest.

